# Hiltonol and Protamine-RNA stimulation induce an immune-activating transcriptome profile in cDC1s

**DOI:** 10.1101/2025.10.13.682171

**Authors:** Georgina Flórez-Grau, Till S. M. Mathan, Mihaela B. Mihaylova, Tom van Oorschot, Gerty Schreibelt, David Sancho, Ignacio Melero, Carl G. Figdor, I. Jolanda M. de Vries, Johannes Textor

**Author notes:** shared first author (Georgina Flórez-Grau, Till S. M. Mathan).

## Abstract

Ex-vivo stimulation of dendritic cells (DCs) is a critical step in DC-based cancer immunotherapies. In humans, conventional type 1 dendritic cells (cDC1s) are a rare myeloid dendritic cell (mDC) subset that express BDCA-3 (CD141). cDC1s promote CD8^+^ T cell cross-priming against tumor antigens and are therefore being explored for use in immunotherapy. We evaluated the impact of ex-vivo stimulation on human peripheral blood cDC1s. In contrast to routine evaluation, which focuses on pre-defined surface maturation markers or soluble factors released from the activated cells, we investigated the impact of stimulation on the transcriptome using both RNA-sequencing (RNA-seq) and microarrays. Specifically, we analyzed the mRNA of cDC1s upon activation with two clinical-grade adjuvants, Hiltonol (poly IC, a TLR3 ligand) and protamine-stabilized RNA (pRNA, a TLR7/8 ligand) compared to unstimulated controls. Both RNA-seq and microarray analysis showed profound and similar effects of both Hiltonol and pRNA on the transcriptome of cDC1s. A gene ontology (GO) analysis suggested that these changes were mainly related to activation and maturation pathways, including induction of type-I interferon (IFN) and interleukin (IL)-12 transcription, while pathways related to adverse effects or cell damage did not appear to be affected. Combination of both reagents did not appear to have a synergistic effect, as the transcriptome changes were similar to those induced by each stimulus alone. Together, our results indicate that both adjuvants have comparable effects on cDC1 maturation within an immunogenic short-term culture as performed in immunotherapy.

## Introduction

DCs are professional antigen-presenting cells that are found in most tissues of the human body.^1^ Upon activation, DCs migrate to the lymph nodes and activate T cells.^2^ Blood-circulating DCs can be divided into two major subsets: mDCs and plasmacytoid dendritic cells (pDCs). The mDCs can be further subdivided into the common conventional type 2 dendritic cell (cDC2) (BDCA1^+^) mDCs and cDC1 (BDCA3^+^) mDCs, which are the rarest subset in the blood.^3,4^ Each DC subset has specific functions. For example, pDCs produce high amounts of type-I IFNs upon sensing viral pathogens via TLR7 or TLR9, while cDC2s sense bacterial pathogens via TLR4 or TLR8 and release IL-12p70 and IL-1β. cDC1s express high levels of TLR3, have been characterized as high IFN-λ and IL-12 producers, and are thought to be highly efficient antigen cross-presenters to CD8 T cells. These properties have led to substantial interest in the use of cDC1s for immunotherapy.^5–12^ Yet, due to their rarity, cDC1s have not been included in clinical trials so far and have not been studied as extensively as the more abundant cDC2s and pDCs.^13,14^

Efficient activation is a crucial step for DC-based cancer immunotherapy.^15–22^ DC activation can be measured using different parameters. Firstly, up-regulation of maturation surface markers like CD40, CD80, CD83, CD86, or HLA-DR as well as chemokine receptors like C-C chemokine receptor type 7 (CCR7) can be measured.^23^ Moreover, to directly test functional maturation, the release of cytokines and chemokines as well as the capacity of DC to prime T cells are measured. All these methods have been used to test different stimuli for DC at good manufacturing practice level.^24,25^ A clear advantage of this analysis is that the data is focused and comparable. However, focusing on selected parameters risks missing important effects that may not be known or expected in advance.

Recently, cDC2s and pDCs have been used for therapeutic vaccination to treat cancer patients, and some favorable clinical outcomes have been observed.^16,19^ In addition, several clinical-grade reagents to induce DC maturation were tested and the complex protamine-RNA (pRNA), which targets TLR8, was suggested to be the most suitable stimulus.^18,26^ Since cDC1s appear promising for future immunotherapies,^27^ we here aimed to identify a suitable stimulus for this DC subset. Because cDC1s are known to express high levels of TLR3,^28^ we tested Hiltonol, as a commercially available clinical grade Poly I:C (PolyICLC) that can trigger TLR3 activation, in addition to the TLR8 ligand pRNA.

To evaluate the suitability of each adjuvant in a comprehensive manner, we applied whole-transcriptome analysis. Specifically, we employed both RNA-seq and microarray measurements to interrogate the DC transcriptome and asked whether these techniques would lead to similar conclusions. Both methods have potential advantages in this context: RNA-seq is more sensitive and therefore increases the chance to observe changes in low-abundant genes, while microarrays can be used with much lower amounts of RNA, which can be a limiting factor for rare cell populations like peripheral blood cDC1s.^29^

## Results

### Microarray and RNA-seq measurements of DC activation lead to similar results

cDC1s (0.7–1.5 million cells) from five donors were isolated and stimulated overnight using each clinical-grade stimulus separately and combined. We used both RNA-seq and microarray-based mRNA analysis on the same samples. Specifically, for 4 out of 5 donors (donors 2-5), there was enough material to perform both RNA-seq and microarray analysis. However, the amount of RNA from donor 1 was insufficient for RNA-seq and therefore only microarray analysis was performed. As a first data exploration step, a principal coordinates analysis (PCoA) was applied to the combined data of all conditions and all donors for both the RNA-seq (**Figure 1A**) and the microarray data (**Figure 1B**). In these plots, the first principal coordinate stratified stimulated and unstimulated cells, whereas the second coordinate appeared to be related to donor differences and no clustering of the different stimuli was observed. Separate PCoA plots per donor (RNA-seq: **Figure 1C**, microarray: **Figure 1D**) also showed the first coordinate to align in each case with stimulation. The combined stimulation was located in between the two individual stimuli in all cases, except for donor 3. For this donor, in both the RNA-seq and the microarray data, the pRNA-stimulated sample did not appear to differ much from the unstimulated samples (highlighted with an arrow in **Figure 1A** and **Figure 1B**). As this outlier was present in both datasets, this indicates that the cells in this sample were not stimulated as expected. The reason for this was not entirely clear – it could have been an issue with this specific sample or genuine variability in how well cDC1 respond to pRNA – and so the best course of action appeared to be to exclude this outlier from the rest of the analysis presented in this paper.

**Figure 1:**
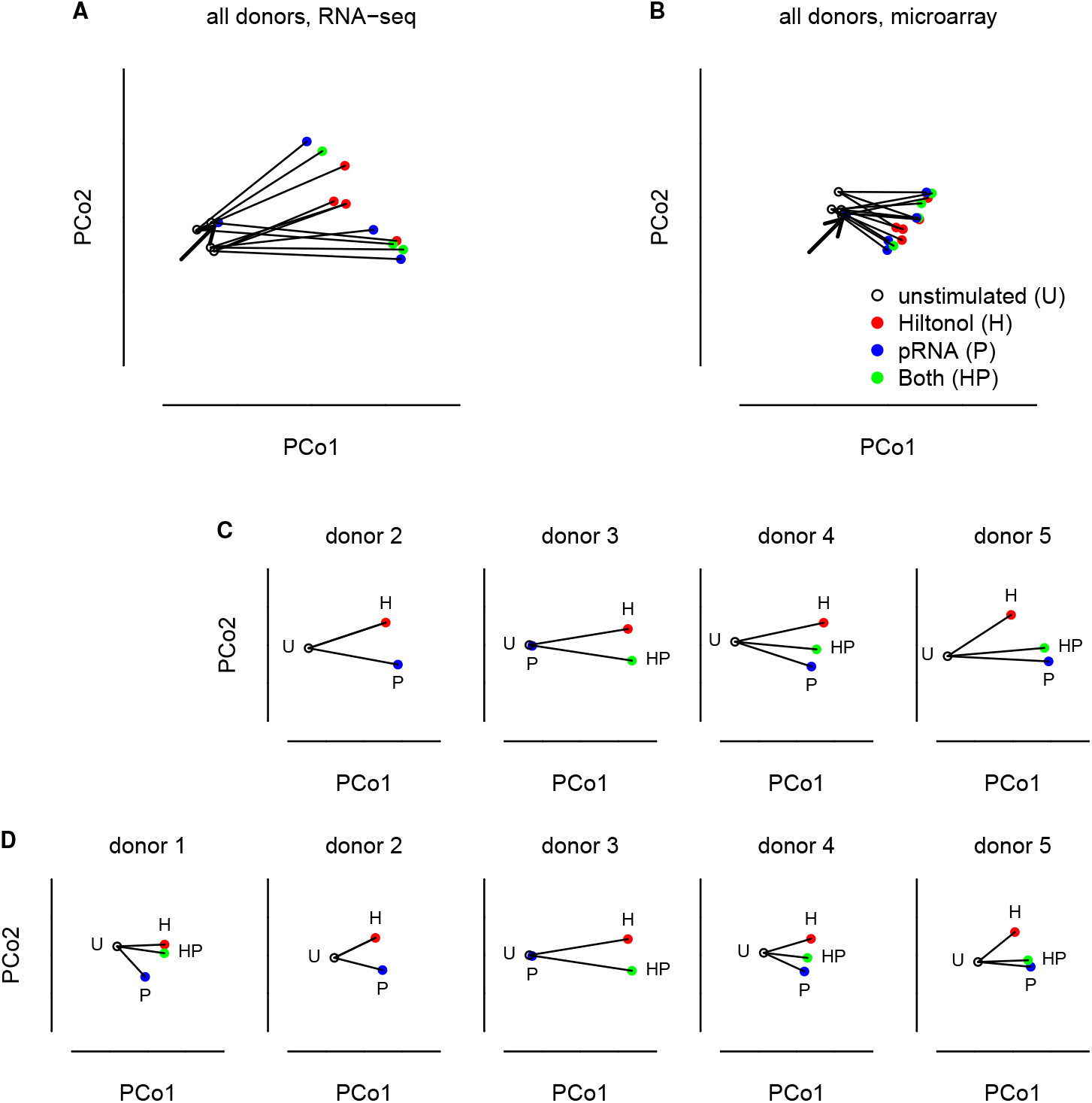
Stimulation of cDC1s is detectable at the transcriptome level. PCoA was performed to globally assess the differences between transcriptomes. Each point represents the transcriptome of one sample and the first and second principal coordinates are plotted. For both the pooled **(A)** RNA-seq and **(B)** microarray datasets, as well as for separate PCoAs for each donor and for (C) RNA-seq and (D) microarray datasets, the first principal coordinate aligned roughly with stimulation. The arrows in (A) and (B) highlight an outlier sample (donor 3) in which the stimulation appears to have failed. This sample was removed from subsequent analysis.

Next, we directly compared the RNA-seq and microarray results for single genes, to see if the conclusions from both analyses are similar. Focusing on the set of genes that showed significant increase or decrease upon stimulation in both datasets, the direction of the change was observed to be the same in all cases (i.e., they were either increased or decreased in both datasets; **Figure 2**). The correlation between the estimated fold changes in the RNA-seq versus microarray data for the selected gene set was 0.9 and 0.91, respectively, largely due to the consistency in the direction of the effect. The fold-change estimated from the RNA-seq datasets was in fact often several orders of magnitude larger than that obtained from the microarray datasets, which was expected due to the higher dynamic range of RNA-seq.^29^ However, this analysis does show that both datasets are expected to yield similar qualitative conclusions, especially in downstream analyses that focus more on the direction of a change than on its magnitude, such as GO analyses.

**Figure 2:**
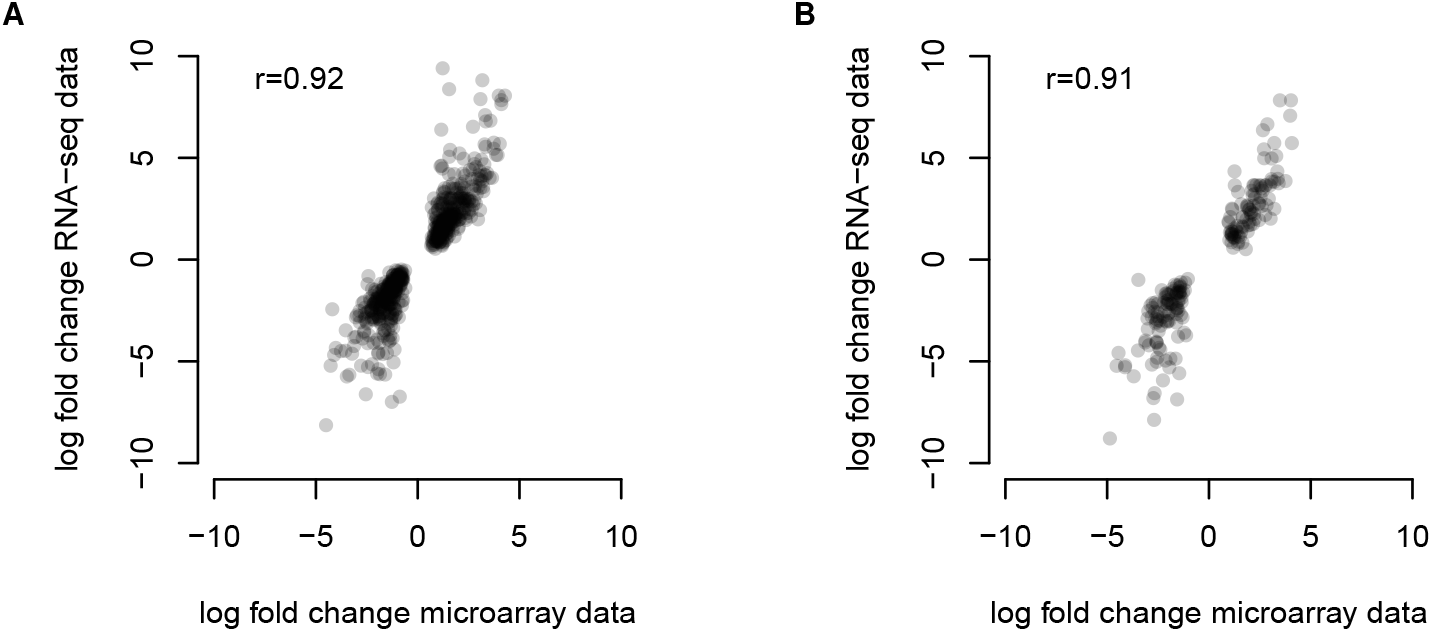
Qualitative agreement between RNA-seq and microarray measurement of stimulation effects. Microarray and RNA-seq fold change estimates for significantly up- or downregulated genes (applying a p-value cut-off of 0.05 after multiple testing correction using the Bonferroni-Holm method) upon stimulation by **(A)** Hiltonol and **(B)** pRNA were plotted against each other. The overall correlation was quantified using Spearman’s rho.

### Hiltonol and pRNA have similar effects on the cDC1 transcriptome

Aiming to identify similarities and differences between the two stimuli, the transcript fold-change values upon Hiltonol or pRNA stimulation were compared to each other for both the RNA-seq (**Figure 3A**) and the microarray (**Figure 3B**) data. This showed that (a) more transcripts were increased upon both stimuli rather than decreased, and that (b) both stimuli had a similar effect; indeed, a differential gene expression analysis based on both datatypes showed no significantly different genes (below a p-value of 0.05 after multiple testing correction) between the two different stimuli (not shown). Focusing on highly upregulated genes (log fold changes >5 for RNA-seq, >2 for microarray) showed, again for both methods, that both stimuli strongly upregulated more transcripts in cDC1 than they downregulated. In summary, both stimuli appear to have similar overall effects on the transcriptome, with both RNA-seq and microarray data supporting this conclusion. For simplicity, we therefore focus on the RNA-seq data in the remainder of this paper.

**Figure 3:**
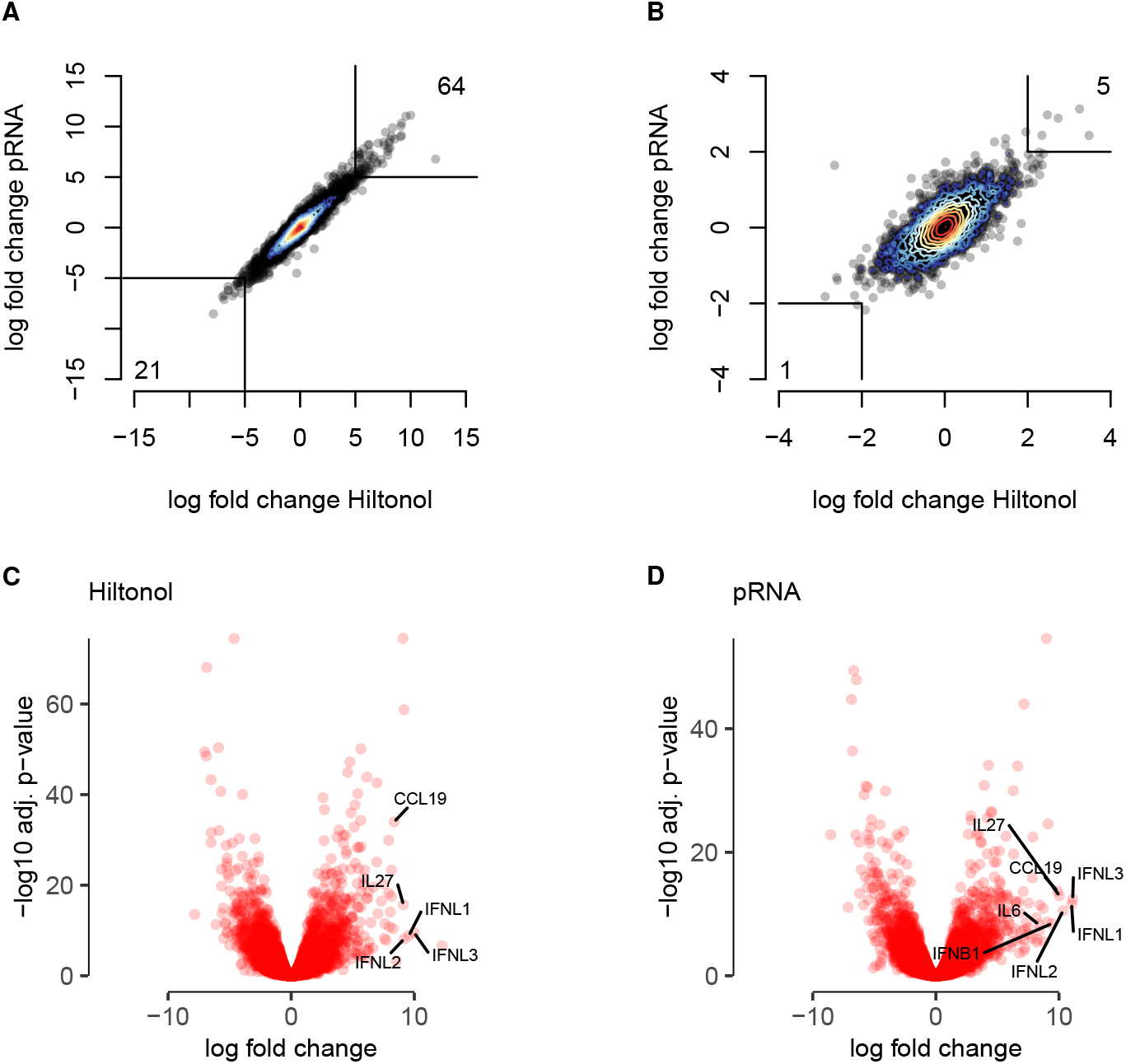
Hiltonol and pRNA stimulation lead to similar changes in the transcriptome of cDC1s. **(A**,**B)** Scatterplots depicting transcript expression changes upon each stimulus as measured by RNA-seq (A) and microarrays (B). Each point represents a transcript. **(C**,**D)** Volcano plots showing the magnitude of transcript abundance changes (x-axis: log fold) together with the statistical significance (y-axis: -log10 p-value). Both stimuli, (C) Hiltonol and (D) pRNA, were compared to the unstimulated sample. A set of transcripts of interest was labeled by name. Fold changes and p-values were determined using edgeR (see Methods). P-values were corrected for multiplicity using the Benjamini-Hochberg method.

To further characterize the most dominant changes upon each stimulus, we generated volcano plots for both stimuli on which we highlighted the expression changes of six cytokines of interest: TNFSF15, IFN-β, IFN-λ, IL-6, IL-27, and CCL19. All were found among the transcripts with the highest expression increase upon stimulation in the RNA-seq data (**Figure 3C,D**). As expected, due to the omission of the outlier sample, the p-values of expression changes for the Hiltonol stimulation were much lower, but the overall log fold changes were very similar.

To identify specific pathways that changed in cDC1 upon stimulation with either Hiltonol or pRNA, a GO term analysis was performed. The more genes belonging to a certain gene cluster change their expression upon stimulation, the higher this gene cluster (i.e. GO term) is ranked. Comparing both stimuli based on the GO term analysis, three overlapping gene clusters among the ten most significant gene clusters of each stimulus were observed. The top differentially expressed GO term upon Hiltonol stimulation was “response to stress”, whereas for pRNA it was “response to virus”. But overall, both stimuli led to an upregulation of similar GO terms, with most terms relating to immune responses or antiviral responses. Importantly, the top 100 affected GO terms contained no pathways related to cell damage or apoptosis (Tables **Figure 1**–**Figure 3**).

### Hiltonol and pRNA stimulation induce a cytokine profile that enhances cross-presentation and cytotoxic immunity

Chemokines, chemokine receptors and interferons are among the most relevant groups of genes for the T cell activating function of DCs.^30–32^ We therefore specifically investigated a range of relevant chemokines, chemokine receptors, interferons, and interleukins to characterize the profile induced by our stimuli in more detail. Given the previous results, we chose not to distinguish between the different stimuli for this analysis.

**Table 1:** GO term analysis comparing unstimulated cells to cells stimulated with Hiltonol.

| term | ontology | N genes | N up/down | $-\log_{10}$ p-value |
| --- | --- | --- | --- | --- |
| protein binding | MF | 14048 | 3386 | 74.8 |
| cytoplasm | CC | 12122 | 3002 | 69.2 |
| molecular function | MF | 18496 | 4100 | 64.5 |
| binding | MF | 16707 | 3809 | 60.1 |
| intracellular anatomical structure | CC | 15487 | 3555 | 47.7 |
| regulation of response to stimulus | BP | 4092 | 1188 | 46.1 |
| positive regulation of biological process | BP | 6375 | 1698 | 43.9 |
| cytosol | CC | 5485 | 1491 | 42.0 |
| response to stress | BP | 4034 | 1143 | 38.5 |
| positive regulation of cellular process | BP | 5813 | 1547 | 38.2 |

**Table 2:** GO term analysis comparing unstimulated cells to cells stimulated with pRNA.

| term | ontology | N genes | N up/down | $-\log_{10}$ p-value |
| --- | --- | --- | --- | --- |
| protein binding | MF | 14048 | 2614 | 55.2 |
| molecular function | MF | 18496 | 3163 | 47.0 |
| binding | MF | 16707 | 2935 | 42.7 |
| cytoplasm | CC | 12122 | 2280 | 41.6 |
| regulation of response to stimulus | BP | 4092 | 932 | 37.4 |
| positive regulation of biological process | BP | 6375 | 1320 | 34.1 |
| response to stress | BP | 4034 | 894 | 30.8 |
| cellular response to chemical stimulus | BP | 2746 | 655 | 30.6 |
| positive regulation of cellular process | BP | 5813 | 1203 | 29.8 |
| cellular response to stimulus | BP | 7599 | 1503 | 29.5 |

**Table 3:** GO term analysis comparing unstimulated cells to cells stimulated with both Hiltonol and pRNA.

| term | ontology | N genes | N up/down | $-\log_{10}$ p-value |
| --- | --- | --- | --- | --- |
| protein binding | MF | 14048 | 3003 | 61.2 |
| molecular function | MF | 18496 | 3652 | 56.4 |
| cytoplasm | CC | 12122 | 2651 | 54.1 |
| binding | MF | 16707 | 3392 | 52.1 |
| regulation of response to stimulus | BP | 4092 | 1064 | 41.4 |
| cellular response to chemical stimulus | BP | 2746 | 761 | 37.8 |
| response to stress | BP | 4034 | 1034 | 37.1 |
| positive regulation of biological process | BP | 6375 | 1502 | 35.9 |
| response to organic substance | BP | 2837 | 771 | 35.1 |
| intracellular anatomical structure | CC | 15487 | 3138 | 34.5 |

We observed that ELR-positive chemokines like CXCL1, CXCL5, and CXCL6 were unaffected or decreased upon stimulation, whereas a consistent increase was seen in the ELR-negative group CXCL9, CXCL10, and CXCL11 (**Figure 4A**). These changes would lead to activation of T cells and NK cells but not neutrophils – suitable for an antitumoral or antiviral immune response. Likewise, the chemokines CCL3, CCL5 and CCL19 – which are involved in the recruitment of T cells and NK cells – all appear to be upregulated.

**Figure 4:**
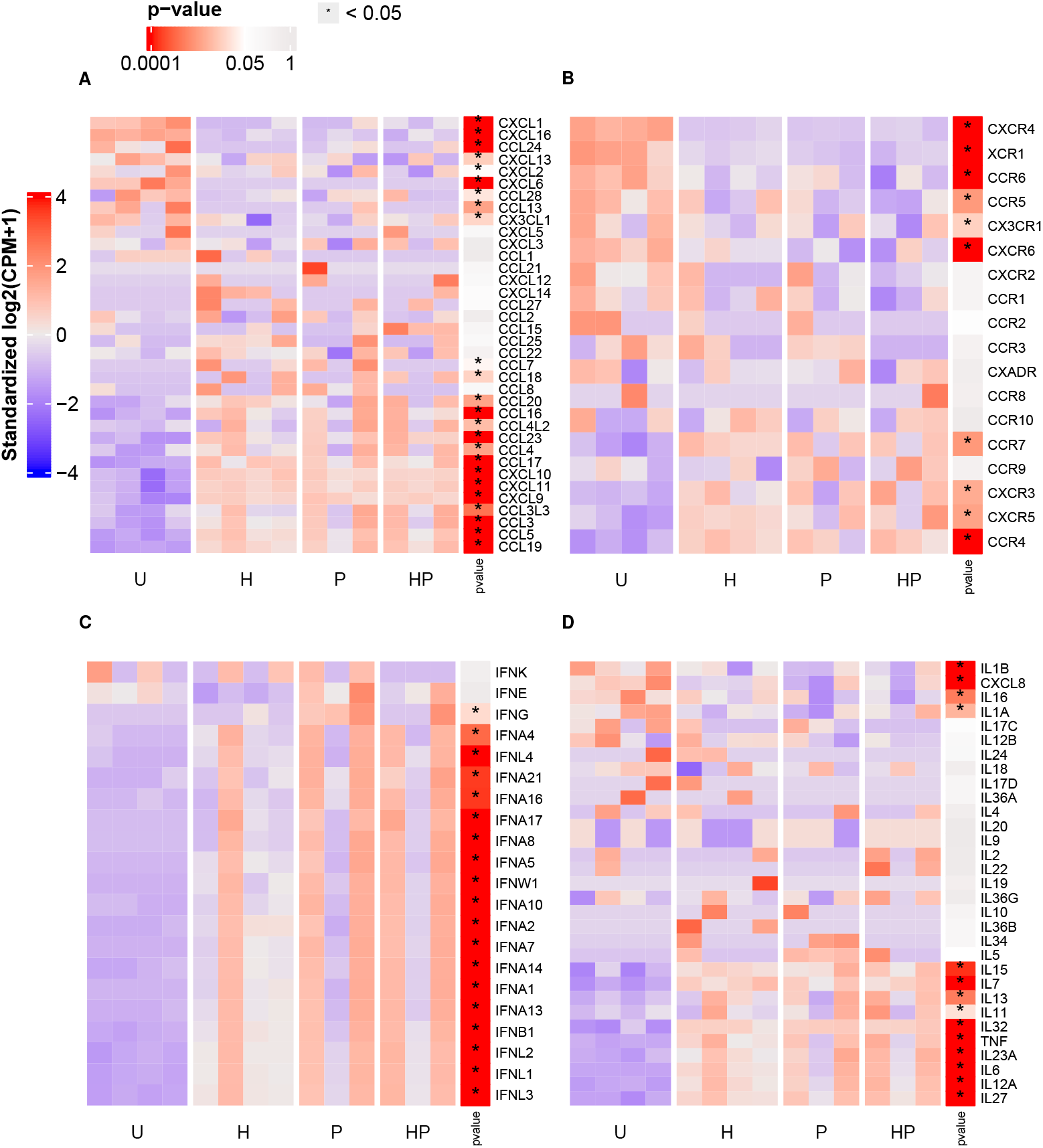
Effect of stimulation on cytokines and cytokine receptors. Heatmaps illustrating the transcript levels in each sample for (A) chemokines, (B) chemokine receptors, (C) interferons, and (D) interleukins. Most transcripts do not appear to be differentially expressed between untreated (U) and treated (H, P, HP) samples. P-values were determined using a likelihood ratio test implemented in edgeR (Methods) where we compared the unstimulated samples to all stimulated samples. P-values were corrected for multiple testing using the Benjamini-Hochberg procedure.

On the chemokine receptor side (**Figure 4B**), an important observation is the decrease in XCR1, which is thought to be important for the recruitment of cDC1 to tumor sites.^32^ This downregulation would prevent activated cDC1s from being retained in peripheral tissues by XCL1/2 gradients, allowing them to instead respond to CCR7 ligands and migrate to lymphoid organs. Consistent with this, there is also a switch from CXCR4 – responding to chemokines expressed in tissues – to CCR4, which has been implicated in DC migration to draining lymph nodes *in vivo*.^33^

The tested interferons showed upregulation across the board (**Figure 4C**), though notably this was less clear for the type II interferon IFNG than for the type I interferons. This was expected due to the stimulation through TLR3 and TLR7/8, which are well-known to mainly lead to production of type I interferons.^30^

In **Figure 4D** we observe that the upregulated interleukins, which include IL6, IL7 and IL12, are all mediators of interactions between DCs and T cells and specifically support CD8 T cell immunity, consistent with the activation of type I interferons seen in **Figure 4C**. A less expected finding is the observed decrease in IL1B and CXCL8 (IL8), since neither cytokine is expected to be produced by immature cDC1s. The initial levels of IL1B were relatively low (3.8 cpm) but those of CXCL8 in fact quite high (676 cpm). For cells that do produce IL1B and CXCL8, such as monocytes or monocyte-derived DCs, we indeed expect this production to be suppressed by type I interferons. It is conceivable that the presence of IL1B and CXCL8 at baseline is a result of culture stress.^34^

Taken together, the observed changes in cytokines and cytokine receptors are consistent with a maturation process where cDC1 would leave peripheral tissues, migrate to draining lymph nodes or other lymphatic sites, and attract and stimulate CD8+ T cells. The detected cytokines would mainly support antiviral and antitumoral immune responses.

### Qualitative comparison of stimuli effects across different DC subsets

The cytokine profile induced by cDC1 stimulation is in some sense reminiscent of pDCs, which are also known to generate primarily type I interferons, migrate to lymph nodes, and stimulate cytotoxic T cells. To corroborate this interpretation, we therefore pooled our cDC1s with RNA-seq data from a previous study on the effects of stimulation on cDC2s and pDCs.^26^ From previous literature^35^ we know that cDC1s and cDC2 are more similar to each other than they are to pDCs; however, given the observations from the cytokine analysis, this might change upon stimulation.

On a joint PCoA plot of all cell populations, the first two principal coordinates aligned with batch effects between the two datasets (PC1) and with a donor difference in the cDC2/pDC dataset (PC2). We therefore show PC3 and PC4 in **Figure 5**. Here, the two conventional DC populations initially appear closer together, but upon stimulation, the cDC1s appear closer to the pDCs. These results lend some additional support to our interpretation of the cytokine changes in **Figure 4** since they were obtained by an analysis of the entire transcriptome rather than a pre-defined set of cytokines and cytokine markers.

**Figure 5:**
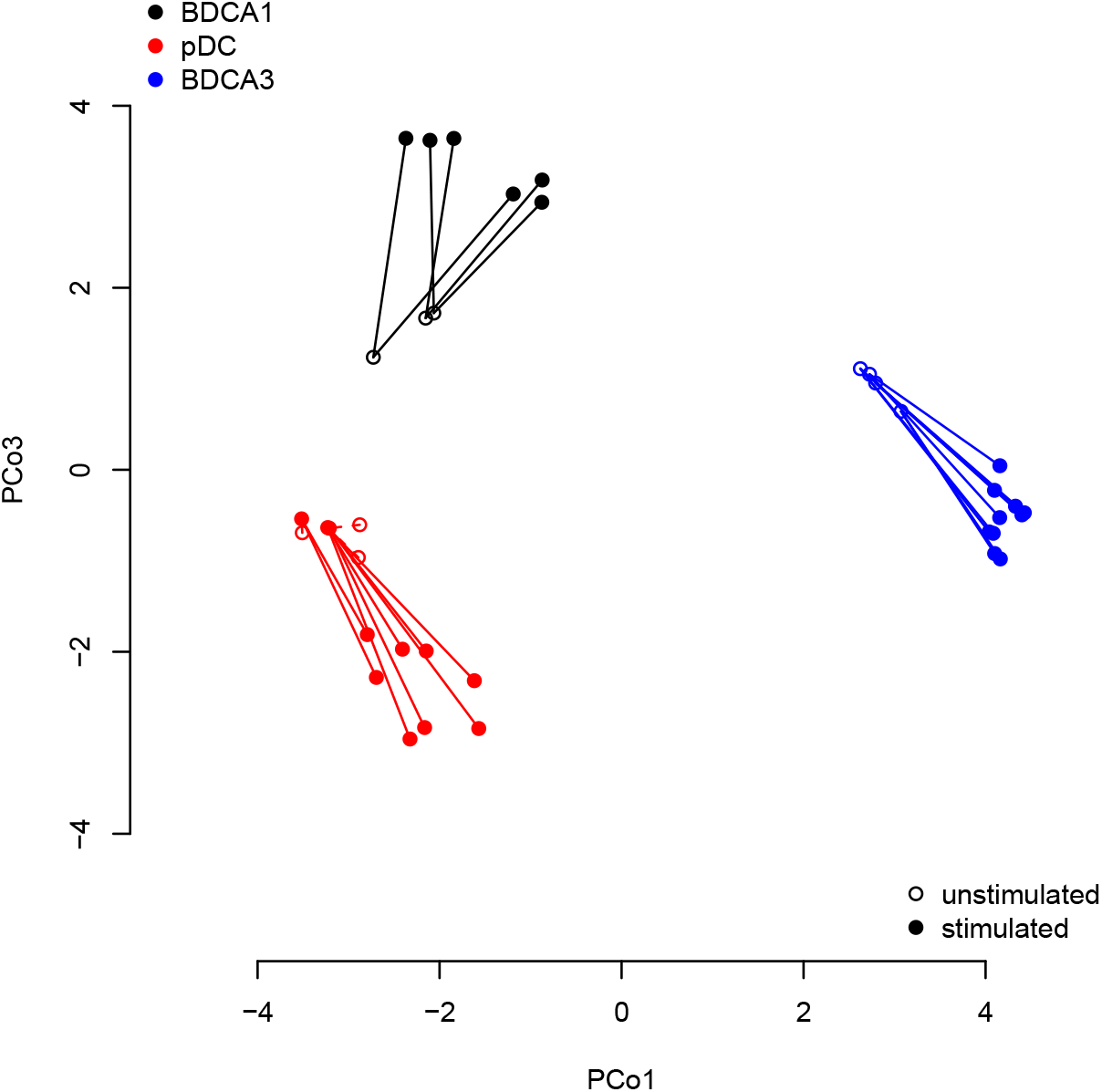
Meta-analysis of stimuli effects on different DC subsets. We pooled our data from this paper with RNA-seq data from an earlier study by Mathan et al.^26^ The pDCs from that study were treated using a mix of IL3 (necessary to keep them alive in culture) and FSME or pRNA. The cDC2s were treated with GM-CSF or pRNA.

### Effects of TLR stimulation on established maturation markers

Finally, we evaluated the transcript levels of established cDC1 maturation markers and therefore investigated the expression of CD80, CD40 and CD86 (**Figure 6A**). The results showed that pRNA led to the strongest increase of CD80 and CD40 transcripts, while CD86 was most increased upon the combination of both stimuli. However, the differences between the stimuli were very minor and not statistically significant. Interestingly, upon stimulation with Hiltonol and pRNA, the C-C chemokine receptor type 7 (CCR7) was upregulated. In contrast, the expression of the MHC class II receptor, HLA-DR, was lower in all stimulated conditions. The combination of the two stimuli had no additional effect on the transcript levels of the maturation markers and the chemokine receptor CCR7.

**Figure 6:**
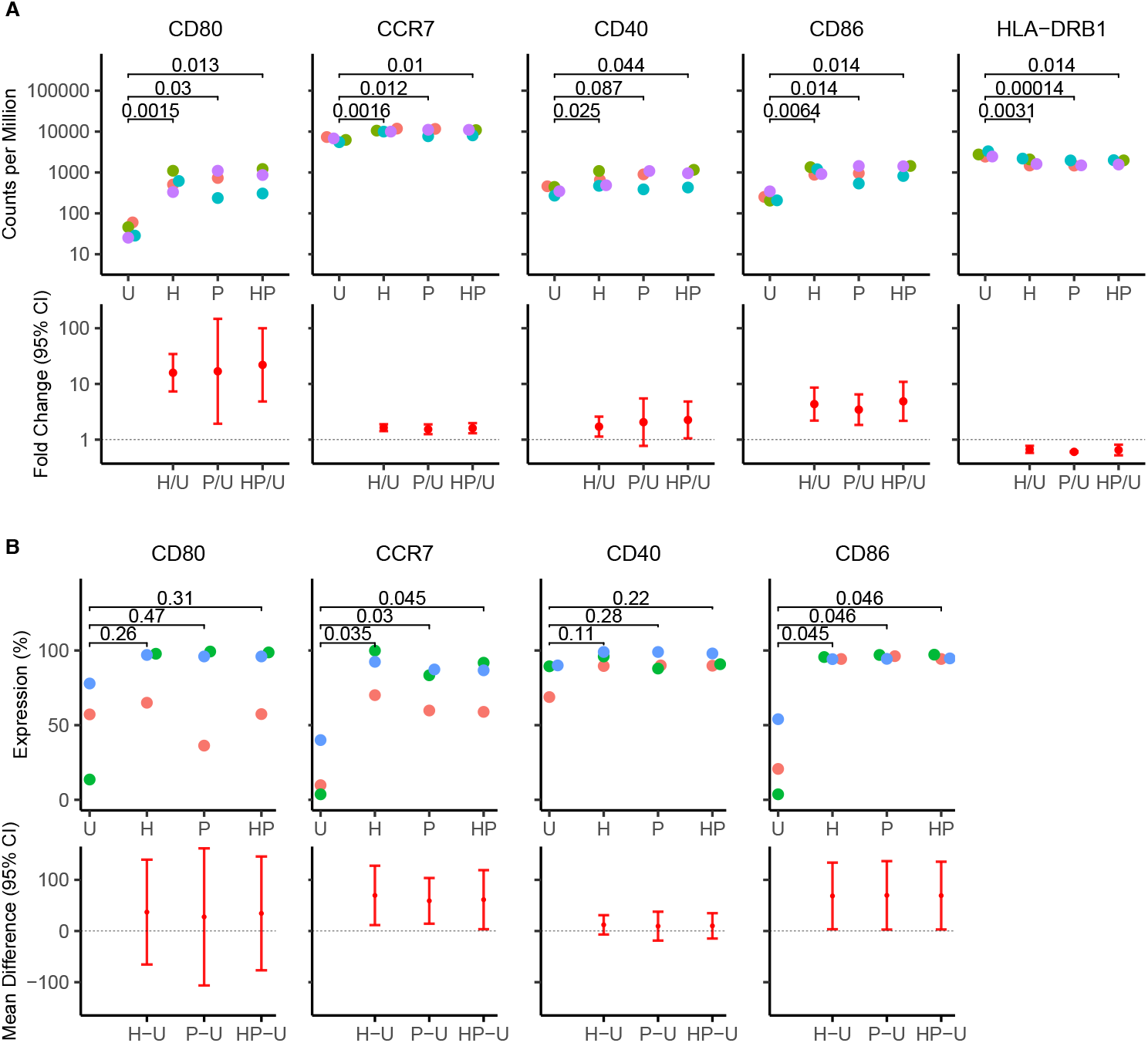
Effect of stimulation on key DC maturation markers. measured by RNA-seq (A) and flow cytometry (B). Each point is one sample, with donors color-coded. The second row of each panel shows the estimated fold changes (A) or differences (B) for each stimulus when compared to the unstimulated sample (error bars: 95% confidence intervals). Different donors were used for the flow cytometry experiments. P-values: paired t tests, not corrected for multiple testing.

In addition, the expression of costimulatory markers was validated at the protein level by flow cytometry analysis (**Figure 6B**). The results of this validation were somewhat mixed – while consistent increases were found in each donor for CCR7 and CD86, the upregulation of CD80 and CD40 was not statistically significant given the substantial variation between donors.

### Effects of TLR stimulation on cytokine release

The release of cytokines that stimulate cytotoxic T cell responses is a particularly important function of DCs in the context of immunotherapy. Even though we have so far not detected major differences between Hiltonol and pRNA, we wanted to investigate this specific aspect of cDC1 maturation in more detail. Therefore, we focused on the four critical cytokines IL6, TNF, IFNA1 and IFNG.

As already seen in **Figure 4**, these cytokines all appeared to be upregulated upon treatment by each stimulus. However, many of these effects are not statistically significant when each stimulus is considered separately, owing to substantial donor variation (**Figure 7A**). In particular, it appears that Hiltonol alone may not trigger upregulation of IFNG transcripts.

**Figure 7:**
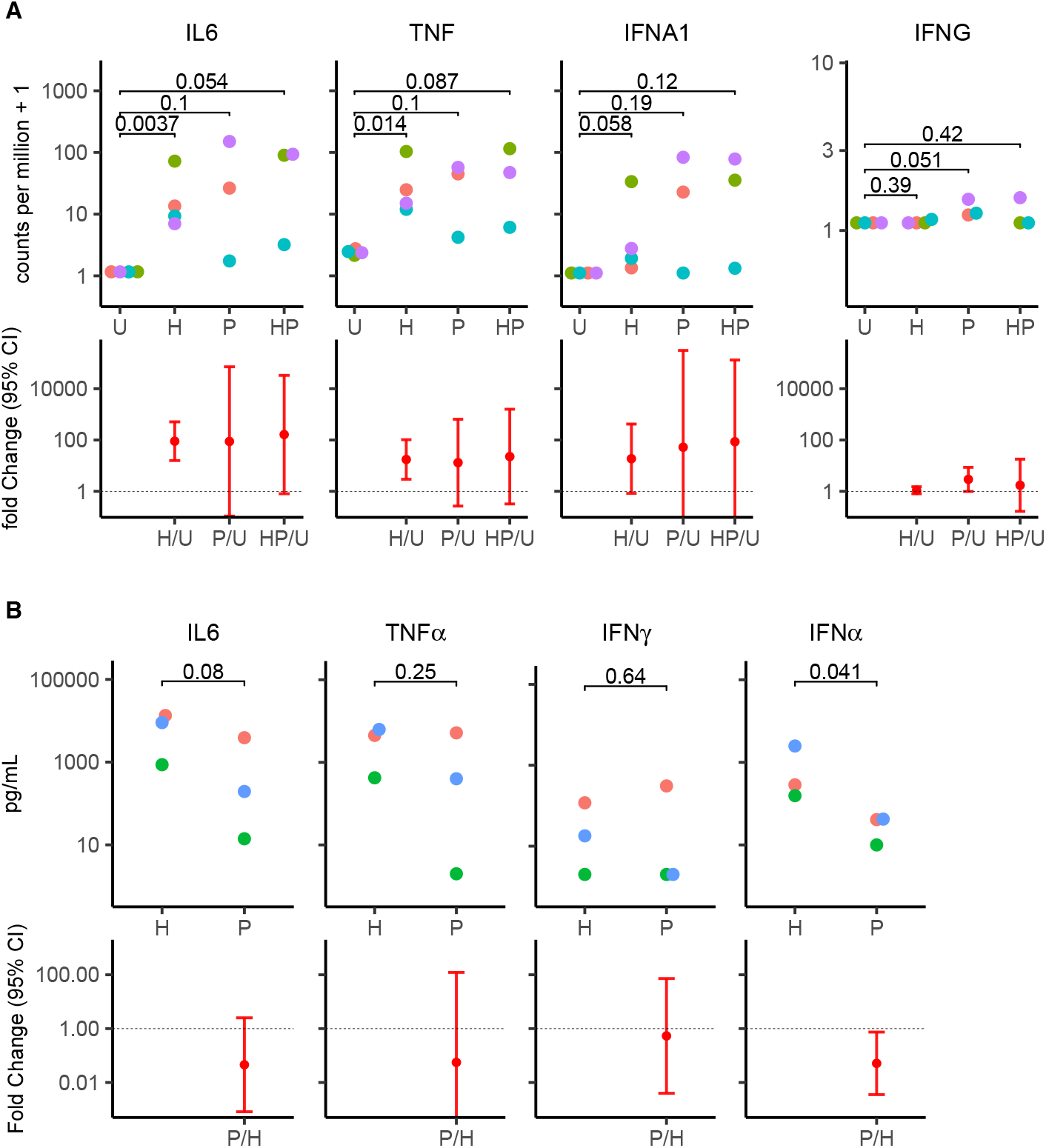
Effects of TLR stimulation on cytokine release. (A) Transcriptome data (raw CPM values underlying the heatmaps shown in Figure 4); (B) ELISA measurements of cytokine release. P-values: paired t tests, not corrected for multiple testing. The second row of each panel shows the estimated fold changes for each stimulus when compared to the unstimulated sample (A) or when comparing Hiltonol to pRNA (B). Error bars: 95% confidence intervals.

To determine whether a difference between the stimuli could be visible on the level of the actually released cytokines, we performed additional ELISA experiments (**Figure 7B**). This analysis did not appear to show major differences between the stimuli, although IFNA production was higher upon Hiltonol treatment, a difference that had not been seen at the transcript level.

Taken together, we did not detect major differences between the stimuli when focusing on four critical cytokines. Larger samples would be needed to answer the question whether such differences exist and could be significant for immunotherapy applications.

## Discussion

In this study, two different clinical-grade TLR stimuli and their effects on cDC1s were compared. The results have demonstrated that both adjuvants present similar effects at the transcriptional level on this subset even though different TLRs were targeted. The analysis was performed by using both RNA-seq and microarrays to analyze the transcriptome and similar conclusions could be drawn suggesting that both stimuli are potent candidates to be used as adjuvants for cDC1 immunotherapeutic applications. Even though pRNA binds most likely TLR7/8 and Hiltonol is a ligand for TLR3, their general effects on cDC1s were strikingly similar, as shown by the strong agreement between the fold-changes estimated for most genes. The combination of both stimuli had a similar effect as either stimulus alone – i.e., we did not find evidence of synergistic effects. Furthermore, our analyses did not indicate any toxic effects of either stimulus on this DC subset, since no GO terms and genes related to adverse responses, e.g., the activation of pathways related to nonsense-mediated decay,^26^ were upregulated or changed upon stimulation. Our findings were in line with a previous study comparing the effect of pRNA and other adjuvants on cDC2 and pDCs, which had suggested a stronger adjuvant potential of pRNA as compared to FSME or GM-CSF.^26^ In addition, our results were in line with an initial study by Sköld *et al*. that characterized the effect of pRNA on DC subsets.^18^

DC maturation state is a critical regulator of immune responses.^31^ Depending on the distinct cytokine patterns released by mature DCs, the polarizing capacity of DCs will differ. In this sense, minor differences regarding cytokine production were observed between the two stimuli (both RNA and protein level). Among the cytokines analyzed, IL-6, TNF-α and Interferons types I and II play an important role in inducing T cell differentiation towards an immunogenic profile.^36^ In this context, a minor difference between pRNA and Hiltonol could be observed by comparing the upregulation of type I/III IFN related genes (Figure 4), which indicated that the combination and predominantly more samples with a pRNA stimulation upregulated type I/III IFN genes. However, these results do not reach statistical significance in the whole-transcriptome analysis. Further functional assays measuring released interferons on the protein level should be performed to further corroborate or disprove this finding. This part of the analysis is crucial to determine which genes related to T cells were activated and therefore predict the maturation status of the cDC1s and the types of interaction with CD8 T cells.

Since the cost of bulk RNA-seq has decreased substantially, the technique has become commercially available and the required amount of RNA has diminished, it has become a standard technique for measuring gene expression, including for some clinical applications.^37–40^ A major advantage of RNA-seq is its ability to quantify and detect novel transcripts, unlike microarray techniques;^41,42^ further, it directly yields RNA sequences, which simplifies data analysis. However, the microarray-based approach used in this study has the advantage that it only requires 20 times less material when compared to the RNA-seq and therefore may be better suited for cell types which are not abundantly available like cDC1s. Importantly, the results of both methods were concordant and the most significant changes were overlapping indicating that microarray measurements might in fact present an attractive alternative for the purpose of evaluating stimulus effects. However, due to its larger dynamic range when measuring transcript abundance, RNA-seq can distinguish more specifically among the lowly expressed genes, which explains why the fold changes estimated from microarray data are often weaker.^29^

In summary, this work provides transcriptome-wide data on how cDC1s react to stimulation by two different clinical-grade stimuli that target different TLRs. A clear advantage of assessing the potential candidate for DC maturation using these approaches is that different information regarding activation status can be obtained with few primary materials, which in the case of cDC1s is crucial due to its lower frequency in human blood. As more parameters than the conventional analysis for maturation (CD80, CD83, CD86, MHC class II and inflammatory cytokines) can be measured, a broader vision of the activation status of the cells is achieved. In consequence, a more accurate selection of the candidate for clinical grade stimuli of DCs can be performed.

## Materials and methods

### Ethics

The study was conducted in accordance with the Declaration of Helsinki. Human peripheral blood samples (buffy coats and apheresis material) were obtained from healthy volunteers at Sanquin (Nijmegen, The Netherlands) after written informed consent. All procedures were approved by the institutional ethics committee of Radboud University Medical Center.

### Cell isolation and culture

For functional assays, DCs were isolated from buffy coats of healthy volunteers (Sanquin, Nijmegen, the Netherlands) after obtaining written informed consent per the Declaration of Helsinki and according to institutional guidelines. For RNA-seq and microarray measurements, cells were obtained from aphaeresis of 5 different donors. Due to the limited cell numbers of donor 1, only 4 donors could be used for RNA-seq. Peripheral blood mononuclear cells (PBMCs) were isolated by using Ficoll density centrifugation (Lymphoprep; Axis-Shield PoC AS, Oslo, Norway). Anti-Human Lineage Cocktail 1 in FITC (LIN1) (BD Bioscience Pharmingen, San Jose, CA) containing antibodies for CD3, CD14, CD16, CD19, CD20, CD56 receptors, together with the anti-FITC conjugated magnetic microbeads of Miltenyi Biotec (Bergisch-Gladbach, Germany) were used to deplete the LIN1^+^ cell fraction, by following manufacturer instructions. Next, cDC1 were further purified by sorting (flowcytometry) using anti-BDCA-3-APC combined with anti-HLA-DR-PE-Vio770 (Miltenyi Biotec) to a purity of 99.9%. DCs were cultured in X-VIVO-15 medium (Lonza, Basel, Switzerland) supplemented with 2% human serum (Sanquin). DCs were stimulated with: pRNA (15 µg/ml) and Hiltonol (10 µg/ml) for 16 hours. Unstimulated controls were cultured overnight in the same medium and conditions as the stimulated cells.

### Protamine-RNA (pRNA) complexes

pRNA complexes were made freshly before addition to the cells. Protamine (protaminehydrochloride MPH 5000 IE/ml; Meda Pharma BV Amstelveen, the Netherlands) was diluted to 0.5 mg/ml in RNase free water and mixed with 2-kbp-long single-stranded mRNA (coding for human gp100 protein).^18^ It was extensively mixed and incubated for 5-10 minutes at room temperature, before being added to the cells.

### RNA sequencing and microarray analysis

cDC1s were isolated as described above and total RNA was extracted using Trizol (Invitrogen, MA, USA), following the standard protocol. The quality control of the isolated RNA (concentration, RIN, 28S/18S and size) was performed with Agilent 2100 Bioanalyzer (Agilent Technologies, Santa Clara, USA). RNA sequencing and read alignment were performed by BGI TECH SOLUTIONS (Hong Kong). Reads were aligned to human genome version 20 (GRCh38). The microarray analysis of the RNA was performed by using the Clariom D assay (Thermo Fisher Scientific).

### Flow cytometry

For validation of maturation markers, purified cDC1s were stained with two antibody panels. Mix 1 included Clec9A–FITC (vio bright), CD40–PE, CD80–PerCP, and CD83–APC. Mix 2 included Clec9A–FITC (vio bright), CCR7–PE, and CD86–APC. Antibodies were titrated according to manufacturer recommendations (Miltenyi Biotec, BD Biosciences) and used at 1:10 dilution. Dead cells were excluded using a viability dye (e788, from Sigma-Aldrich). Samples were acquired on a BD FACSVerse (BD Biosciences) and analyzed using FlowJo v10. Fluorescence minus one (FMO) controls were included for gating. Data are presented as mean fluorescence intensity (MFI) or percentage of positive cells relative to unstimulated controls.

### ELISA

Selected cytokines (IL-6, TNF, IFN-α, and IFN-γ) were quantified in cell culture supernatants using commercial ELISA kits (R&D Systems), according to the manufacturers’ instructions. Briefly, 96-well plates pre-coated with capture antibody were incubated with diluted supernatants and standards, followed by biotinylated detection antibodies and streptavidin-HRP. The reaction was developed using TMB substrate and stopped with 2 N H_2_SO_4_. Absorbance was measured at 450 nm with wavelength correction at 570 nm using an iMark Microplate Reader (Bio-Rad). Cytokine concentrations were calculated from standard curves included on each plate.

### Statistical analysis

Data were analysed using the R environment for statistical computing, version 4.2.1.^43^ It contained raw RNA counts, as well as microarray data and we were interested in studying changes in gene expression in the three experimental conditions - treatment with Hiltonol (H), treatment with pRNA (P), and combined treatment with both Hiltonol and pRNA (HP), compared to the untreated condition (U).

The package edgeR,^44–46^ in BiocManager,^47^ was used for whole-transcriptome principal coordinates analysis, differential gene expression analysis, and GO term analysis. For the principal coordinates analysis we turned the table of raw counts into a DGEList() object and then scaled it, using the calcNormFactors() function with the default method (TMM). We then used the plotMDS() function to generate separate plots for the RNA-seq and microarray data (Fig. 1). This allowed us to identify an outlier - the treatment with pRNA had not worked on one of the donors and this donor was excluded from the rest of the analysis.

We calculated differential gene expression between the untreated and each one of the treatment conditions. We used the raw counts and created a design matrix using model.matrix(), with treatment condition as a factor. We then estimated the dispersion parameter using estimateDisp(), using the design matrix and default parameters. Differential expression was determined by fitting a generalized linear model using glmFit(), and significance was determined using the likelihood ratio test provided by the glmLRT() function. In our GO term analysis, we used all differentially expressed genes for each of the three comparison pairs (H-U, P-U, and HP-U). We used a Benjamini-Hochberg correction for multiple comparisons adjustment and a significance cutoff of 0.05 for the adjusted p-values.

We generated correlation plots for logFC values of genes that were differentially expressed for both the Hiltonol and the pRNA treatments, for both the RNA-seq and microarray data (Fig. 3a,3b). We used a custom function, plt_corr(), which uses ggplot2.^48^ In addition, we generated volcano plots for the adjusted p-values for the log-fold change under the conditions H, P, and HP, compared to U, using the RNA-seq data. These plots were generated with a custom function, plt_adj_p(), which uses ggplot2.

We also visualised gene expression changes for some cytokines, chemokines, and type-I interferons, across the four conditions (Fig. 4a, b, c, d). For this purpose, we used the ComplexHeatmap^49,50^ package, and, more precisely, the Heatmap() function. For the visualisation we scaled the logCPM data to values in (0,1). Clustering was switched off. The p-values shown are obtained from conducting a repeated-measures ANOVA on the non-scaled logCPM values for the different conditions. The p-values are adjusted for multiple comparisons using the Benjamini-Hochberg correction.

Furthermore, we visualised the expression of a set of important maturation markers, namely, CD80, CCR7, CD40, CD86, and HLA-DRB1 (Fig. 5a, b, c, d, e), using the custom gene_swarm_diff_plot() function. We calculated the mean differences in expression between H, P, HP, and the untreated condition U, using a paired two-tailed t-test. The p-values displayed have not been adjusted for multiple comparisons. Similar plots were also used for Fig.7a, b, but in 7a we visualised log-fold changes and used another custom function, gene_swarm_fold_plot().

It is important to note that the design of **Figure 6** and **Figure 7** was heavily influenced by the package dabestr.^51^ Our implementation uses two-tailed t-tests instead of bootstrapping given the sample size. We used a two-tailed t-test, instead of bootstrapping, as our sample size was very low. In addition, we added a plot of the effect sizes of the log-fold changes.

## Data availability

RNA-seq and microarray data have been deposited in NCBI’s Gene Expression Omnibus^69^ and are accessible through GEO Series accession number GSE239899 (https://www.ncbi.nlm.nih.gov/geo/query/acc.cgi?acc=GSE239899).

## Code availability

Our analysis code will be shared publicly when this paper has undergone peer review and the analyses are final. In the meantime, we are happy to share the code with anyone interested upon request.

**Table 4:**
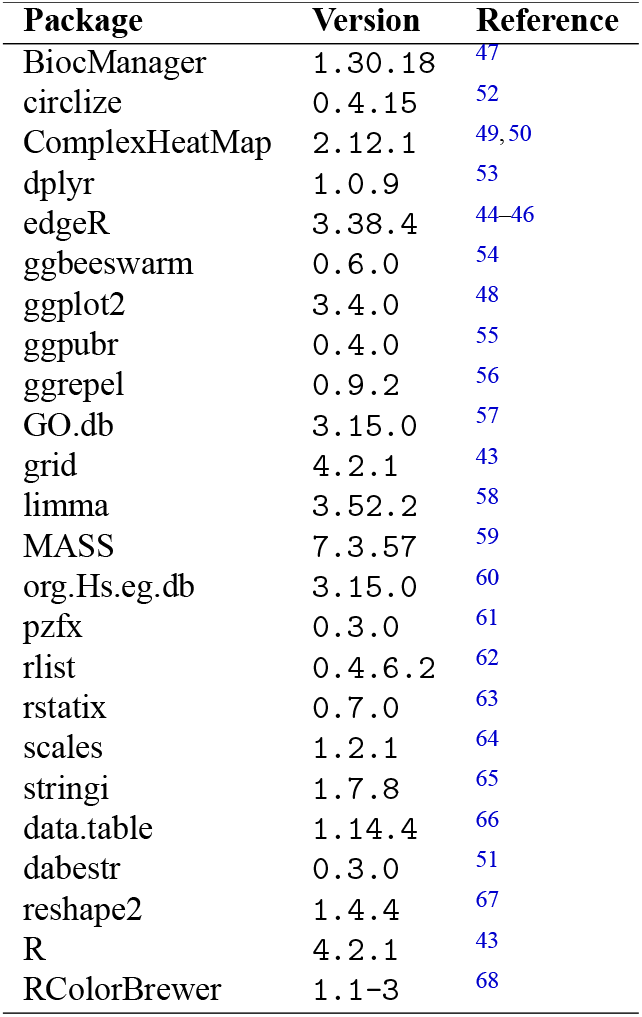
R Packages with Citations (Alphabetically)

## Acknowledgements

We thank Thermo Fisher Scientific for advice on experimental design and for processing the Clariom D samples to enable us to perform the RNA-Seq comparisons.

## Author contributions statement

JdV, GS, DS, IM, TM, and CGF jointly designed the research. TM, TvO, and GFG performed experiments. JT, MBM and TM analyzed the data. TM, JT, GFG and MBM wrote the manuscript. All authors critically revised the manuscript for important intellectual content.

## List of abbreviations

cDC1: conventional type 1 dendritic cell
cDC2: conventional type 2 dendritic cell
DC: dendritic cell
GO: gene ontology
IFN: interferon
IL: interleukin
mDC: myeloid dendritic cell
pDC: plasmacytoid dendritic cell
PCoA: principal coordinates analysis

